# NFE2L1-mediated proteasome function protects from ferroptosis

**DOI:** 10.1101/2021.12.12.472274

**Authors:** Stefan Kotschi, Anna Jung, Nienke Willemsen, Anahita Ofoghi, Bettina Proneth, Marcus Conrad, Alexander Bartelt

**Author notes:** Correspondence: Univ.-Prof. Dr. rer. nat. Alexander Bartelt, Professor of Cardiovascular Metabolism, Institute for Cardiovascular Prevention, IPEK, Ludwig-Maximilians-University Munich, Klinikum der Universität München AöR, Pettenkoferstraße 9, 80336 München.

## Abstract

**Objective:** Ferroptosis continues to emerge as a novel modality of cell death with important therapeutic implications for a variety of diseases, most notably cancer and degenerative diseases. While susceptibility, initiation, and execution of ferroptosis have been linked to reprogramming of cellular lipid metabolism, imbalances in iron-redox homeostasis, and aberrant mitochondrial respiration, the detailed mechanisms of ferroptosis are still insufficiently well understood.

**Methods and Results:** Here we show that diminished proteasome function is a new mechanistic feature of ferroptosis. The transcription factor nuclear factor erythroid-2, like-1 (NFE2L1) protects from ferroptosis by sustaining proteasomal activity. In cellular systems, loss of NFE2L1 reduced cellular viability after the induction of both chemically and genetically induced ferroptosis, which was linked to the regulation of proteasomal activity under these conditions. Importantly, this was reproduced in a Sedaghatian-type Spondylometaphyseal Dysplasia (SSMD) patient-derived cell line carrying mutated glutathione peroxidase-4 (GPX4), a critical regulator of ferroptosis. Also, reduced proteasomal activity was associated with ferroptosis in *Gpx4*-deficient mice. In a mouse model for genetic *Nfe2l1* deficiency, we observed brown adipose tissue (BAT) involution, hyperubiquitination of ferroptosis regulators, including the GPX4 pathway, and other hallmarks of ferroptosis.

**Conclusion:** Our data highlight the relevance of the NFE2L1-proteasome pathway in ferroptosis. Manipulation of NFE2L1 activity might enhance ferroptosis-inducing cancer therapies as well as protect from aberrant ferroptosis in neurodegeneration, general metabolism, and beyond.

**Graphical abstract:** 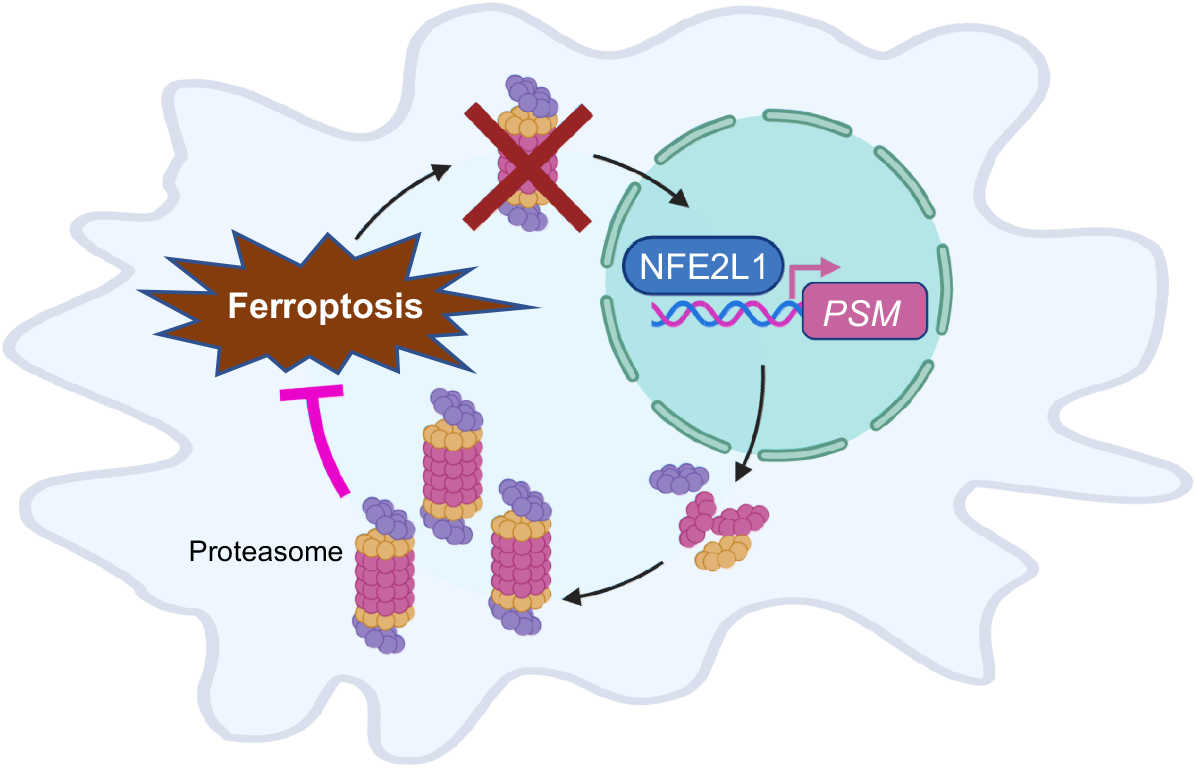

**Highlights:**

- Proteasome function is diminished during ferroptosis
- NFE2L1-mediated proteasomal activity protects from ferroptosis
- The ubiquitination of the GPX4-glutathione pathway is implicated in *Nfe2l1* deficiency
- NFE2L1 deficiency in brown fat is associated with hallmarks of ferroptosis

## 1. Introduction

Ferroptosis is an iron-dependent, lipid peroxidation-mediated non-apoptotic form of cell death [1]. Ferroptosis has only been recognized relatively recently as a non-apoptotic entity of cell death and remains poorly understood [2, 3]. Deciphering the mechanisms responsible for initiation or prevention of ferroptosis has become a thriving field of research as ferroptosis has been implicated in various forms of cancer and degenerative diseases [3, 4]. The peroxidation of polyunsaturated fatty acid residues esterified in phospholipids has been established as the key event leading to ferroptosis. Lipid peroxidation occurs in the presence of unbound iron by the Fenton reaction [5]. Both lipid antioxidants such as α-tocopherol or ferrostatin-1 and iron chelators can effectively prevent ferroptosis in cells [2, 6].

Glutathione-peroxidase 4 (GPX4) is a key enzyme for preventing lipid peroxidation and ferroptosis [7, 8]. GPX4 is a selenoprotein that efficiently catalyzes the reduction of lipid hydroperoxides in cellular membranes at the expense of glutathione (GSH) or other protein thiols [9, 10]. Pharmacological inhibition of GPX4 by the small molecule RSL3 leads to cell death in cultured cells [7]. In mice, germline deletion of the *Gpx4* gene is lethal in utero, and adult mice with an inducible deletion of *Gpx4* die within 2 weeks from acute kidney failure [8, 9]. Human mutations in *GPX4* are extremely rare, but a few patients have been identified displaying Sedaghatian-type Spondylometaphyseal Dysplasia (SSMD), a neonatal disease with skeletal abnormalities, central nervous system malformations and early mortality [11]. The antioxidant capacity of GPX4 is dependent on the availability of reduced GSH or sufficient cysteine in case of limiting GSH levels in cells. Likewise, the synthesis of GSH is dependent on cellular cystine imported by the cystine-glutamate antiporter system Xc- (encoded by *SLC7A11*). Inhibiting system Xc-by the compound erastin ultimately causes cysteine starvation, GSH depletion, and ferroptosis [2]. Furthermore, degradation of system Xc- and GPX4 is carried out by the ubiquitin-proteasome system (UPS), and studies have shown that deubiquitinase and proteasome inhibitors impact on GPX4 levels and ferroptosis in cancer cells [12, 13]. However, whether UPS is causally involved in ferroptosis is unclear.

Damaged, unwanted, or obsolete proteins are targeted for degradation by a complex sequence of posttranslational events of the UPS, resulting in the polyubiquitination of proteins, which are ultimately deubiquitinated and processed by the proteasome. The proteasome is a macromolecular ATP-dependent protease complex consisting of proteasomal subunits (encoded by *PSM* genes) and regulatory parts. To prevent proteotoxicity, endoplasmic reticulum (ER) stress, and cellular dysfunction, global ubiquitination levels need to be matched by proper proteasomal activity [14]. The transcription factor NFE2L1 has emerged as a key regulator of the adaptive component of the UPS, as it enhances the expression of *PSM* genes when proteostasis is challenged [14]. NFE2L1 undergoes complex posttranslational regulation [15]. Partly embedded in the membrane of the ER, NFE2L1 requires deglycosylation by N-glycanase-1 and proteolytic cleavage, resulting in the liberation of an approximately 95 kDa fragment containing the DNA-binding domain. This active fragment is being constantly degraded and is virtually absent, but when proteasome function is compromised, e.g., in the presence of proteasome inhibitors, the 95 kDa fragment enables the recovery of proteasomal activity in a bounce-back mechanism by stimulating the gene expression of proteasomal subunits [16].

Global *Nfe2l1* deficiency results in embryonic lethality [17], but conditional models have shed light on the biological relevance of NFE2L1-mediated proteasomal activity. We have previously shown that NFE2L1 is a critical regulator of cellular adaptation and stress resistance under certain physiologic and pathological conditions. Interestingly, cold adaptation in mice and humans requires an adaptive increase in proteasomal protein quality control and this mechanism of thermogenic adaptation in brown adipocytes is mediated by NFE2L1 [18]. Furthermore, NFE2L1 has been shown to be involved in the regulation of lipid metabolism by directly sensing ER cholesterol and mitigating the deleterious effects of excess cholesterol [19]. While some of the downstream pathologies involve ER stress, lipotoxicity, and inflammation, the exact proximal outcome of NFE2L1 dysfunction remains unclear. The interplay between NFE2L1-mediated proteasomal function and ferroptosis has not been investigated under these conditions, but previous studies have indicated a potential link between GPX4, UPS, and ferroptosis. Here, we investigate the role of NFE2L1 in ferroptosis and its implications for proteasomal function in cell systems, transgenic animal mouse models, and in a SSMD patient-derived human cell line carrying a mutated GPX4.

## 2. Material and Methods

### 2.1. Cell culture and treatments

Cells were cultured in growth medium DMEM GlutaMax (Gibco), 10% fetal bovine serum (FBS, Sigma) and 1 % penicillin-streptomycin (PS, Sigma). Upon confluence, immortalized brown adipocytes (imBAT) cells were differentiated to mature adipocytes by the addition of 1 μM rosiglitazone (Cayman Chemicals), 1 μM T3 (Sigma-Aldrich), 850 nM human insulin (Sigma), 500 nM IBMX (Sigma), 1 μM dexamethasone (Sigma) and 125 nM indomethacin (Sigma) for 48 h, after which the medium was changed to growth medium containing only rosiglitazone, T3 and insulin with renewing this medium every subsequent 48 h. Experiments were carried out on the day 6 of the differentiation protocol. The human fibrosarcoma cell line HT-1080 was cultured in DMEM high glucose (Sigma) enriched with 10 % FBS and 1 % PS. The peradipocyte cell line WT1 was cultured in DMEM high glucose (Sigma) enriched with 10 % FBS and 1 % PS. Chemical induction of ferroptosis was performed using 10 μM erastin (Selleck Chem), 1 μM RSL3 (Selleck Chem) and 10 μM FIN56 (Selleck Chem). Proteasome inhibitors were used as 100 nM bortezomib (Selleck Chem) and 100 nM epoxomicin (Millipore). Ferroptosis was inhibited using 10 μM ferrostatin-1 (Selleck Chem). For experiments with WT1 and HT-1080 cells, the compounds were added 6 h after reverse transfection into the transfection medium. Treatment of imBAT cells was added 24 h after transfection, while changing the medium to differentiation medium. Knockdown experiments were performed using SMARTpool siRNA (Dharmacon). Reverse transfection was performed using Lipofectamine™ RNAiMAX transfection reagent (Thermo Fisher) and siRNA for *Nfe2l1/NFE2L1* and *Gpx4* at a concentration of 30 nM. For experiments, 50,000 cells were seeded in a 24-well format, 10,000 cells in a 96-well format, and 500,000 cells in a 6-well format, respectively. imBAT cells were transfected at day 4 of differentiation. Primary fibroblasts from a patient with SSMD caused by homozygous mutation c.647 G>A in exon 6 of GPX4 (GPX4 mut) and his healthy father (GPX4 WT) were provided by Sanath K. Ramesh (curegpx4.org). SSMD cells were cultured in DMEM GlutaMax supplemented with 10 % FBS and 1 % PS. As GPX4 mut cells do not proliferate and undergo cell death under normal conditions, 10 μM ferrostatin-1 was added to the medium to prevent lipid peroxidation and ferroptosis. For assessment of cell viability, 15,000 cells per well from both cell lines were transfected into 96-well-plates, as specified below. For proteasomal activity assays in SSMD cells, 300,000 cells were seeded per well of a 6-well-plate. During transfection/seeding supplementation of ferrostatin-1 was ended. The medium was changed 24 h after transfection, and 10 μM ferrostatin-1 was supplemented again where specified. AquaBluer (MultiTarget Pharmaceuticals) was used to assess cell viability. 20 h after treatments, medium was changed and replaced with 1:100 AquaBluer in phenol red-free DMEM GlutaMax (Thermo Fisher) medium and incubated for 4 h at 37 °C. Fluorescence was measured at 540ex/590em using a Spark Multimode Microplate Reader (Tecan).

### 2.2. Mice and treatments

All animal experiments were performed according to procedures approved by the animal welfare committees of the government of Upper Bavaria, Germany (ROB-55.2-2532.Vet_02-20-32, ROB-55.2-2532.Vet_02-20-51) and performed in compliance with German Animal Welfare Laws. Mice carrying floxed *Nfe2l1* alleles and Ucp1-Cre have been previously described [18]. Animals were housed in individually ventilated cages at room temperature with a 12 h light-dark cycle. All mouse housing and husbandry occurred on standard chow diet (Ssniff). For histology and proteomics, we housed at 4 °C for 7 days for cold acclimatization. To measure non-shivering thermogenesis, we injected norepinephrine (NE, Sigma, 1 mg/kg in 0.9 % NaCl) in a Sable Systems Promethion Indirect Calorimetry System. 10-12 weeks old Gpx4^fl/fl^ ROSA26^CreERT2^ or Gpx4^fl/fl^ control mice were injected twice i.p. with 100 μL of a tamoxifen solution (Sigma, 20 mg/mL) dissolved in Mygliol. After 7 days, the mice were euthanized, and tissues were harvested.

### 2.3. Native Page and proteasomal activity

The protocol for in-gel proteasome activity assay and subsequent immunoblotting was described previously [20]. In short, cells were lysed in OK Lysis buffer (50 mM Tris/HCl (pH 7.5), 2 mM DTT, 5 mM MgCl_2_, 10 % glycerol, 2 mM ATP, 0.05 % Digitonin), containing phosphate inhibitor (PhosStop, Roche). The suspensions were kept on ice for 20 min and then centrifuged twice. Protein concentration in the supernatant was determined using Bradford assay (Bio-RAD) according to the manufacturer’s protocol. In a NuPAGE 3-8 % Tris-Acetate gel (NOVEX Life Technologies), 15 μg protein was loaded with 5 x loading buffer (0.05 % Bromophenol Blue, 53.5 % glycerol, 250 mM Tris). The gel was run at a constant voltage of 150 V for 4 h. Afterwards, the gel was incubated in an activity buffer (50 mM Tris, 1 mM MgCl_2_, 1 mM DTT) with 0.05 mM chymotrypsin-like substrate Suc-Leu-Leu-Val-Tyr-AMC (Bachem) for 30 min at 37 °C. The fluorescent signal was measured using ChemiDoc MP (Bio-RAD). The gel was then incubated in solubilization buffer (2 % SDS, 66 mM Na_2_CO_3_, 1.5 % β-mercaptoethanol) for 15 min to prepare the samples for blotting. Through tank transfer, the samples were transferred to a PVDF membrane on 40 mA overnight. The PVDF membrane was blocked in ROTI-block and incubated in primary antibody (1:1000) overnight at 8 °C and subsequently with secondary antibody (1:10,000) for 3 h at room temperature (see Supplementary Table 1). To assess proteasomal activity in cells and tissues, the chemotrypsin-like, trypsin-like and caspase-like activity were measured using the Proteasome Activity Fluorometric Assay (UBP Bio) as specified in the manufacturer’s manual and previously described [18]. Normalization to protein concentration was carried out with Bio-Rad Protein Assay Kit II in the same samples.

### 2.4. NFE2L1 reporter assay

HEK293a cells stably expressing firefly luciferase driven by upstream activator sequence (UAS) promotor and the chimeric NFE2L1, in which DNA-binding domain was replaced with the Gal4 UAS-targeting sequence, were treated as described previously [19]. This assay measures nuclear translocation and transactivation of luciferase by the NFE2L1-UAS binding to its promoter. 25,000 cells were plated per well of a 96-well plate in DMEM GlutaMax (Thermo Fisher) containing 10 % fetal bovine serum (Sigma) and 1% penicillin-streptomycin (Sigma). After treatment with the specified compounds cells were harvested and luciferase emission measured using Dual-Glo^®^ Luciferase Assay System (Promega) according to the manufacturer’s instructions. Normalization to DNA content was carried out using Quant-iT™ PicoGreen™ dsDNA Assay Kit (Thermo Fisher).

### 2.5. Protein extraction and immunoblotting

Protein extraction and immunoblotting were performed as previously described [21]. Briefly, cells or tissues were lysed using a TissueLyser II (3 min, 30 Hz; Qiagen) in RIPA buffer (150 mM NaCl (Merck), 5 mM EDTA (Merck), 50 mM Tris pH 8 (Merck), 1 % IGEPAL^®^ CA-630 (Sigma), 0.5 % sodium deoxycholate (Sigma-Aldrich), 0.1 % SDS (Roth) containing 1 μM protease inhibitors (Sigma). Protein concentration was determined by Pierce BCA Protein Assay (Thermo Fisher Scientific) according to the manufacturer’s instructions. Lysate protein concentrations were adjusted in Bolt™ LDS Sample buffer (Thermo Fisher) containing 5 % β-mercaptoethanol (Sigma). SDS-PAGE was performed using Bolt™ 4–12 % Bis-Tris gels (Thermo Fisher) in Bolt™ MOPS SDS running buffer. Proteins were transferred onto a 0.2 μm PVDF membrane (Bio-Rad) using the Trans-Blot^®^ Turbo™ system (Bio-Rad) according to the manufacturer’s instructions. The membrane was stained in Ponceau S (Sigma-Aldrich for determination of equal protein concentration and blocked in Roti-Block (Roth) for 1 h at room temperature. Primary antibodies (see Supplementary Table 1) were incubated overnight at 4 °C. Blots were washed 3 x for 10 min with TBS-T (200 mM Tris (Merck), 1.36 mM NaCl (Merck), 0.1 % Tween 20 (Sigma)) and secondary antibodies (see Supplementary Table 1) applied for 1 h at room temperature. After subsequent washing in TBS-T (3 x for 10 min) membranes were developed using SuperSignal West Pico PLUS Chemiluminescent Substrate (Thermo Fisher) in a Chemidoc imager (Bio-Rad).

### 2.6. Proteomics and analysis

The raw proteasome and ubiquitome data were described previously [18]. Differential expression analysis was performed using DeSeq2 [22] and KEGG pathway analysis and GO term analysis was achieved with Clusterprofiler [23] in R.

### 2.7. RNA extraction, cDNA synthesis and qPCR

RNA extraction was performed using the NucleoSpin^®^ RNA kit (Macherey-Nagel) as specified by the manufacturer. For RNA extraction from tissue, samples were lyzed in TRIzol™ (Thermo Fisher) using a TissueLyser II (3 min, 30 Hz; Qiagen). After vigorous mixing with chloroform at a 1:4 v/v ratio (chloroform:TRIzol), samples were centrifuged and the supernatant transferred onto the purification columns of the NucleoSpin^®^ RNA kit. All further steps were executed as specified by the manufacturer. cDNA was synthesized with Maxima™ H Master Mix 5 x (Thermo Fisher) using 500 ng of RNA. Gene expression was evaluated by qPCR using PowerUp™ SYBR Green Master Mix according to the manufacturer’s instructions. Primers are listed in Supplementary Table 1. Expression was normalized to *Tbp/TBP* levels.

### 2.8. Statistics

Data are shown as the mean ± standard error of the mean (SEM). For comparisons of two groups, we used student’s T-test, for three and more groups we used 1-way ANOVA with Bonferroni post-hoc test as indicated in the figure legends. For comparing two variables we used 2-way ANOVA followed by Bonferroni post-hoc test. Analysis was performed using R and/or GraphPad Prism. A *P*-value < 0.05 was considered significant, as indicated by asterisks in the figures or as indicated otherwise.

## 3. Results

### 3.1. Proteasome function is inhibited during ferroptosis

The proteasome is a critical part of the UPS and a major mediator of cellular stress resistance. This involves both the specific and targeted degradation of key molecules as well as the global disposal of ubiquitinated proteins to maintain proper proteostasis and suppress cellular stress. Therefore, we asked if ferroptosis is linked to the modulation of proteasomal function. Shortterm incubation with the chemical inducers of ferroptosis erastin, RSL3 and FIN56 [24] led to lower levels of proteasomal activity at non-lethal concentrations (Fig. 1A). Interestingly, longer incubation times led to a bounce back of proteasomal activity (Fig. 1A). In line with the compromised proteasomal activity during the early stages of ferroptosis, the expression of key regulatory proteasomal subunits was lower with erastin and RSL3 treatment (Fig. 1B). To study the assembly of the proteasome complex during ferroptosis, we performed native PAGE, during which the tertiary structure of the proteasome is preserved [20]. Confirming the findings of the SDS-PAGE, both 30S and 20S proteasomal activity and expression were lower after the induction of ferroptosis by erastin and RSL3 (Fig. 1C), indicating acute impairment of proteasomal function during the induction of ferroptosis. Next, we investigated proteasomal function in models of “genetic” induction of ferroptosis. In the global inducible knockout model of *Gpx4*, proteasomal activity was markedly lower compared to controls in various tissues (Fig. 1D, E). To translate these findings to a human pathological ferroptosis model, we used primary fibroblasts from a patient carrying an autosomal recessive homozygous mutation in *GPX4* causing SSMD. These cells require cultivation in the presence of lipid antioxidants, for instance ferrostatin-1, because otherwise they rapidly succumb to ferroptosis. In line with our previous findings, after withdrawal of ferrostatin-1 proteasomal activity was markedly lower in the SSMD cells compared to control fibroblasts from a healthy donor, which were not affected by ferrostatin-1 (Fig. 1F). However, proteasome gene expression, exemplified by *PSMA1* and *PSMB1* mRNA levels (Fig. 1G), was higher in the SSMD cells compared to control fibroblasts, and this effect was suppressed by ferrostatin-1. These results show that proteasome function is compromised upon the induction of ferroptosis. Furthermore, this is also associated with an adaptive response by the cells.

**Figure 1:**
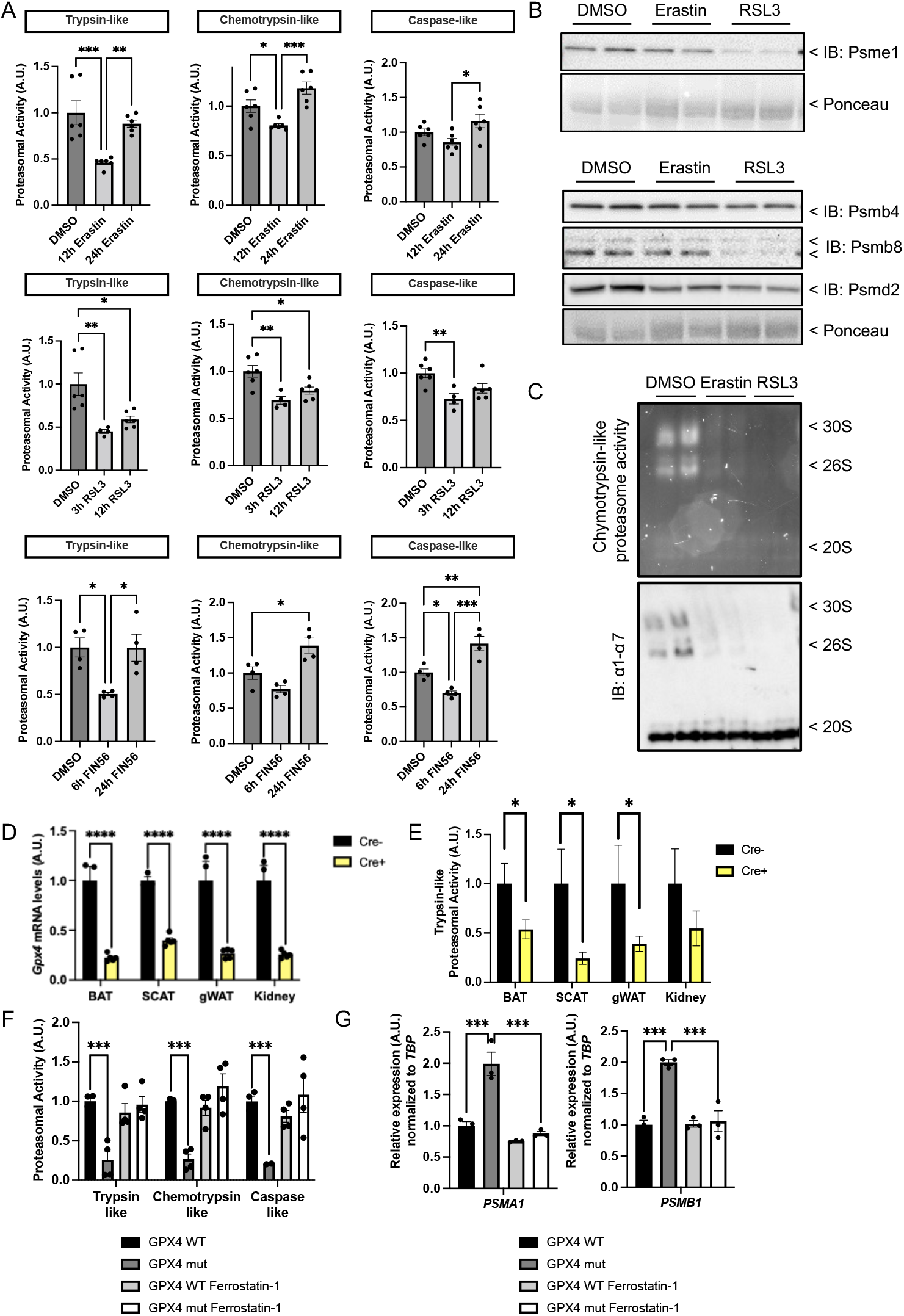
Loss of proteasomal function is a hallmark of ferroptosis. **(A)** Proteasomal activity in imBAT cells treated with 10 μM erastin for 12 h and 24 h, 1 μM RSL3 for 3 h and 12 h, or 10 μM FIN56 for 6 h and 24 h (*n* = 4-6 replicates) **(B)** Immunoblot of proteasome subunits in WT1 cells after treatment with 10 μM erastin or 1 μM RSL3 for 6 h **(C)** Native Page of WT1 cells treated with 10 μM erastin and 1 μM RSL3 for 6 h with in-gel measurement of proteasomal activity and immunoblot of the α1-7 subunits (representative images from 3 experiments) **(D,E)** Gpx4 mRNA levels and proteasomal activity in tissues of *Gpx4*^fl/fl^ ROSA26^CreERT2^ mice (*n* = 3-5 biol. replicates) **(F)** Proteasomal activity in SSMD cells (GPX4 mut) and healthy controls (GPX4 WT) after 24 h of treatment with 10 μM ferrostatin-1 (*n* = 4 replicates). **(G)** Proteasome subunit mRNA levels (*n* = 3 replicates). Statistical significance: \**P*_adj_ < 0.05 (0.1 in E), \*\**P*_adj_ < 0.01, \*\*\**P*_adj_ < 0.001, \*\*\*\**P*_adj_ < 0.001 by 2-way ANOVA followed by Tukey’s test **(A,D-F)**. BAT: brown adipose tissue, SCAT: subcutaneous adipose tissue, gWAT: gonadal white adipose tissue

### 3.2. Nfe2l1 is activated during ferroptosis

The adaptive restoration of proteasomal activity, e.g., in the presence of proteasome inhibitors, is mediated by NFE2L1 activation [16]. Interestingly, induction of ferroptosis with RSL3 or FIN56, but not erastin led to higher NFE2L1 protein levels (Fig. 2A). These changes were not observed on *NFE2L1* mRNA levels (Fig. 2B), as erastin led to higher and RSL3 to lower levels. The discrepancies between mRNA and protein levels point to a post-transcriptional regulation of NFE2L1 activity. To investigate if these changes in protein levels also reflect changes in the transcriptional activity of NFE2L1, we used an established reporter cell line expressing firefly luciferase under control of the UAS promoter and NFE2L1 fused to a UAS-targeting DNA-binding domain [19]. Both RSL3 and FIN56 treatment led to higher NFE2L1 activity in a time-dependent manner, to a similar extent as the proteasomal inhibitor bortezomib, a known activator of the NFE2L1 pathway (Fig. 2C, D). However, treatment with erastin led to a decrease in NFE2L1 activity (Fig. 2E). Taken together, induction of ferroptosis by RSL3 and FIN56, but not with erastin increases protein levels and activity of NFE2L1.

**Figure 2:**
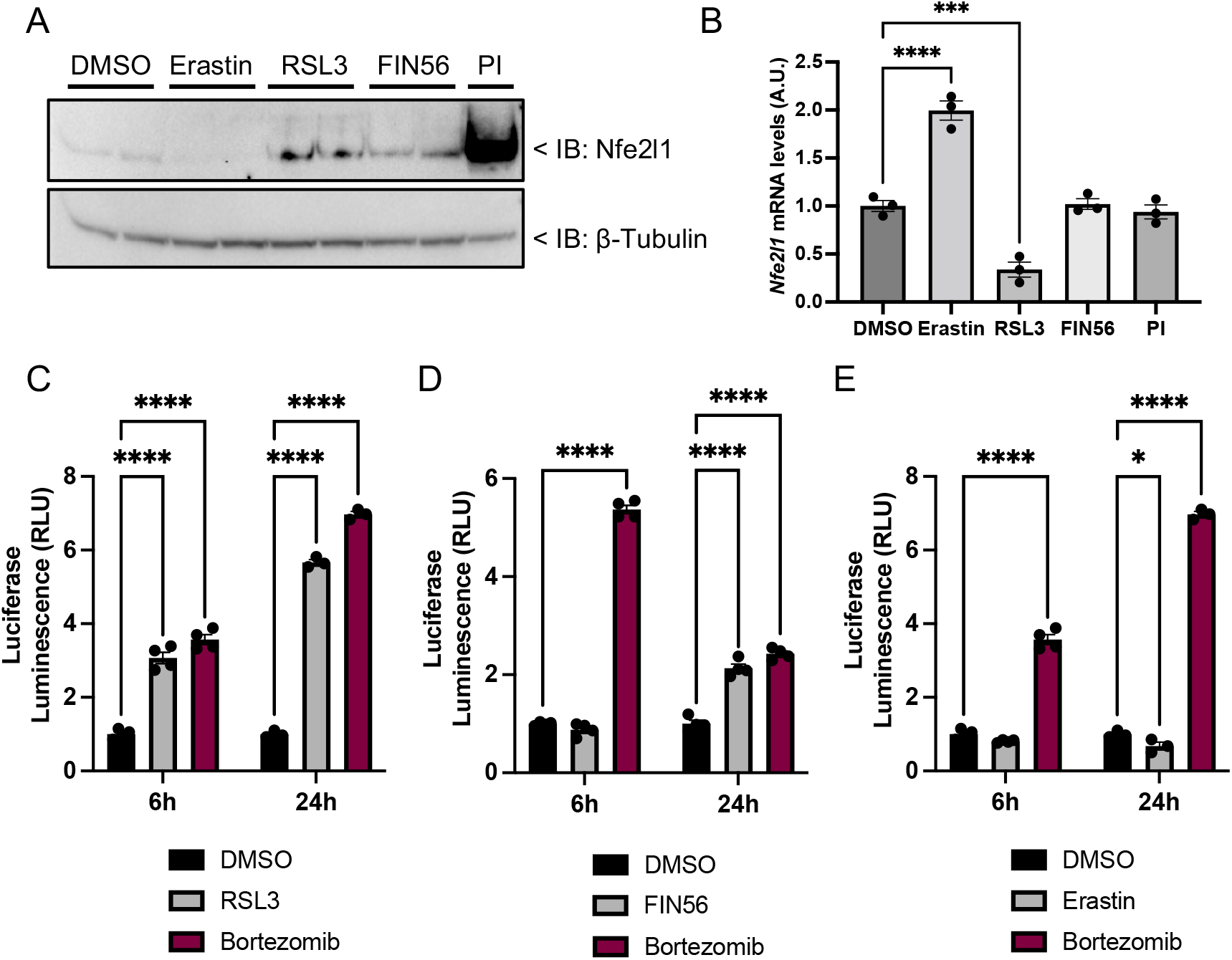
Nfe2l1 is activated during ferroptosis with RSL3 and FIN56. **(A)** Immunoblot of Nfe2l1 and β-tubulin in WT1 cells treated with 10 μM erastin, 1 μM RSL3, 50 μM FIN56, or 100 nM proteasome inhibitor (PI) bortezomib for 8 h **(B)** mRNA levels of *Nfe2l1* in imBAT cells treated with 10 μM erastin, 1 μM RSL3, 10 μM FIN56 and 100 nM PI bortezomib for 12 h normalized to the expression of the housekeeping gene *Tbp* (*n* = 3 replicates). **(C-E)** NFE2L1 luciferase reporter assay in HEK293a cells treated treatment with (A) 1 μM RSL3, (B) 50 μM FIN56, (C) 10 μM erastin and 100 nM bortezomib for 6 h and 24 h (*n* = 3-4 replicates) Statistical significance: \**P*_adj_ < 0.05, \*\**P*_adj_ < 0.01, \*\*\**P*_adj_ < 0.001, \*\*\*\**P*_adj_ < 0.0001 by 2-way ANOVA followed by Tukey’s test **(B-E)**.

### 3.3. Nfe2l1 protects from ferroptosis in vitro

To further evaluate the link between NFE2L1 and ferroptosis, we used a cellular system, in which we acutely targeted NFE2L1 expression by siRNA. To confirm the efficacy of the knockdown, cells were also treated with the proteasome inhibitor bortezomib, which usually increases protein levels of the cleaved 95 kDa form of NFE2L1. In both untreated and bortezomib conditions, NFE2L1 was silenced on the protein level (Fig. 3A). Also, as expected, silencing of NFE2L1 led to lower levels of *PSM* gene expression (Fig. 3B). Next, we investigated cell viability during pharmacological induction of ferroptosis using erastin, RSL3 and FIN56. Significantly lower cell viability was found after treatment with the ferroptosis-inducing compounds RSL3 and FIN56 when NFE2L1 was silenced (Fig. 3C). As inhibition of GPX4 leads to ferroptosis, we also evaluated “genetic” induction of ferroptosis by siRNA-mediated knockdown of GPX4 in conjunction with additional knockdown of NFE2L1. 24 h after siRNA transfection, only cells lacking both GPX4 and NFE2L1 showed lower cell viability (Fig. 3D). Additionally, we used SSMD cells, in which silencing of NFE2L1 markedly lowered viability in comparison to control cells (Fig. 3E, F). This phenotype was rescued by the addition of ferrostatin-1 24 h after transfection, restoring viability (Fig. 3F). These results show that NFE2L1 protects from ferroptosis. Interestingly, while proteasome inhibitors itself do not induce ferroptosis, we wondered whether the protective role of NFE2L1 in ferroptosis might relate to its adaptive regulation of proteasome function. To address this question, we treated SSMD and control cells with knockdown of NFE2L1 with bortezomib in the presence and absence of ferrostatin-1 (Fig. 3G). While these treatments did not reduce viability of control cells, SSMD cells showed lower viability, which was rescued by ferrostatin-1 treatment. This effect was markedly more pronounced when the SSMD cells were treated with bortezomib in the absence of NFE2L1, which also was rescued by ferrostatin-1. Taken together, these results demonstrate that NFE2L1-mediated proteasome function protects from ferroptosis.

**Figure 3:**
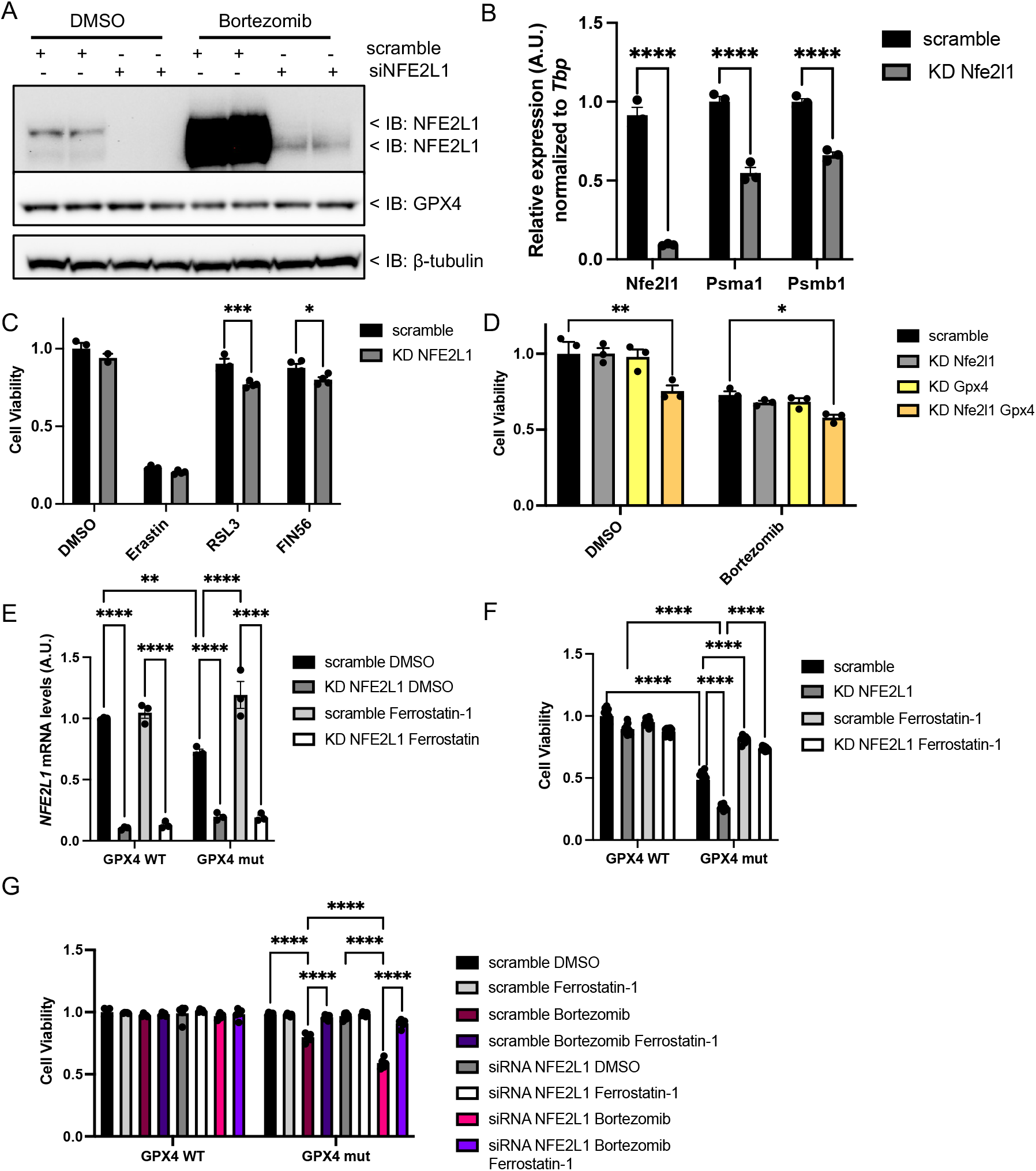
Loss of NFE2L1 increases susceptibility to ferroptosis. **(A)** Immunoblot of Nfe2l1 and β-Tubulin in HT-1080 cells treated with NFE2L1-targeting siRNA and 100 nM bortezomib (*n* = 2 replicates) **(B)** mRNA levels of *Nfe2l1, Psma1* and *Psmb1* in imBAT cells during siRNA-mediated knockdown of *Nfe2l1* normalized to the expression of the housekeeping gene *Tbp* (*n* = 3 replicates) **(C)** Cell viability of HT-1080 cells treated with 10 μM erastin, 1 μM RSL3 or 10 μM FIN56 for 24 h (*n* = 4 replicates) **(D)** Cell viability of WT1 cells 24 h after siRNA-mediated inhibition of Nfe2l1 and/or Gpx4 (*n* = 3 replicates) **(E)** *NFE2L1* mRNA levels in SSMD cells (GPX4 mut) and healthy controls (GPX4 WT) after siRNA-mediated knockdown of *NFE2L1*. Cells were cultured without ferrostatin-1 after transfection, 24 h later 10 μM ferrostatin-1 was supplemented as indicated (*n* = 3 replicates) **(F)** Cell viability of SSMD cells and healthy controls 60h after siRNA-mediated knockdown of *NFE2L1*. (*n* = 10 replicates) **(G)** Cell viability of SSMD and control cells during siRNA-mediated knockdown of NFE2L1 and incubation with 100 nm bortezomib and/or 10 μM ferrostatin-1 for 12 h. Statistical significance: \**P*_adj_ < 0.05, \*\**P*_adj_ < 0.01, \*\*\**P*_adj_ < 0.001, \*\*\*\**P*_adj_ < 0.0001 by 2-way ANOVA followed by Tukey’s test **(B-G)**.

### 3.4. Hallmarks of ferroptosis in a mouse model deficient in Nfe2l1

While these data in cells indicate that NFE2L1 suppresses ferroptosis, the biological significance of this process remains unknown. Interestingly, we observed that proteasome function in brown adipose tissue (BAT) of mice lacking *Gpx4* was reduced (Fig. 1D, E), indicating that there might be a link between ferroptosis, proteasome, and NFE2L1 in BAT. To test this hypothesis, we used a recently created transgenic mouse model of conditional *Nfe2l1* deficiency, which has helped us to discover new physiological roles for NFE2L1 in vivo [18 19]. BAT is a mammalian-specific organ dedicated to non-shivering thermogenesis, which is sustained by NFE2L1-mediated proteasome function [18]. Thus, NFE2L1 maintains proteostasis under high metabolic activity, and in the absence of NFE2L1, brown adipocytes develop malfunctional UPS and ER stress [18]. To determine if this NFE2L1-dependent stress response in BAT was associated with ferroptosis, we analyzed the proteome of BAT isolated from mice lacking *Nfe2l1* and controls [18]. Gene ontology analysis revealed that the major changes in the proteome were related to mitochondrial function and many aspects of metabolism (Fig. 4A, B). Interestingly, among these was a large panel of genes linked to ferroptosis, including known critical ferroptosis regulators such as GPX4 [10] and acyl-CoA Synthetase Long Chain Family Member 4 (ACSL4) [25], that were expressed at higher levels in BAT isolated from *Nfe2l1*-deficient animals compared to controls (Fig. 4C). To investigate the impact of insufficient proteasomal activity in this model, we analyzed ubiquitome data, which were generated by immunoprecipitation of tryptic ubiquitin remnant motifs [18]. In the absence of Nfe2l1, BAT displayed a higher ubiquitination status compared to controls (Fig. 4D). As GPX4 activity and its supply with GSH is a critical protective mechanism that has been linked to UPS [12, 13], we investigated ubiquitination status of GPX4, and the enzymes required for the synthesis of GSH (Fig. 4E, F). Glutamate-cysteine ligase (GCL) is the first rate-limiting enzyme of GSH synthesis and consists of two subunits, a heavy catalytic subunit GCLC and a light regulatory subunit GCLM. We identified lysine residues representing ubiquitination sites of GCLC and GLCM that were significantly enriched in BAT isolated from *Nfe2l1*-deficient animals (Fig. 4E, F), indicating hyperubiquitination. Also, we found that glutathione synthetase (GSS), the second and final enzyme for GSH synthesis displayed a marked increase in ubiquitination based on higher representation of K186 in BAT isolated from *Nfe2l1*-deficient animals (Fig. 4E, F). Finally, we identified two ubiquitination sites, K148 and K168, in GPX4 that were significantly enriched in BAT isolated from *Nfe2l1*-deficient animals, indicating a hyperubiquitination Gpx4 (Fig. 4E, F). Interestingly, when exposed to cold BAT deficient in *Nfe2l1* undergoes a complex whitening process (Fig. 4G) that diminishes oxygen consumption in response to norepinephrine, indicating blunted thermogenic capacity (Fig. 4H). In summary, these data highlight the role of NFE2L1-mediated proteasome function for BAT. Based on our global analysis, the absence of NFE2L1 is characterized by hallmarks of ferroptosis, which is linked to the ubiquitination status of the GSH-GPX4 pathway in vivo.

**Figure 4:**
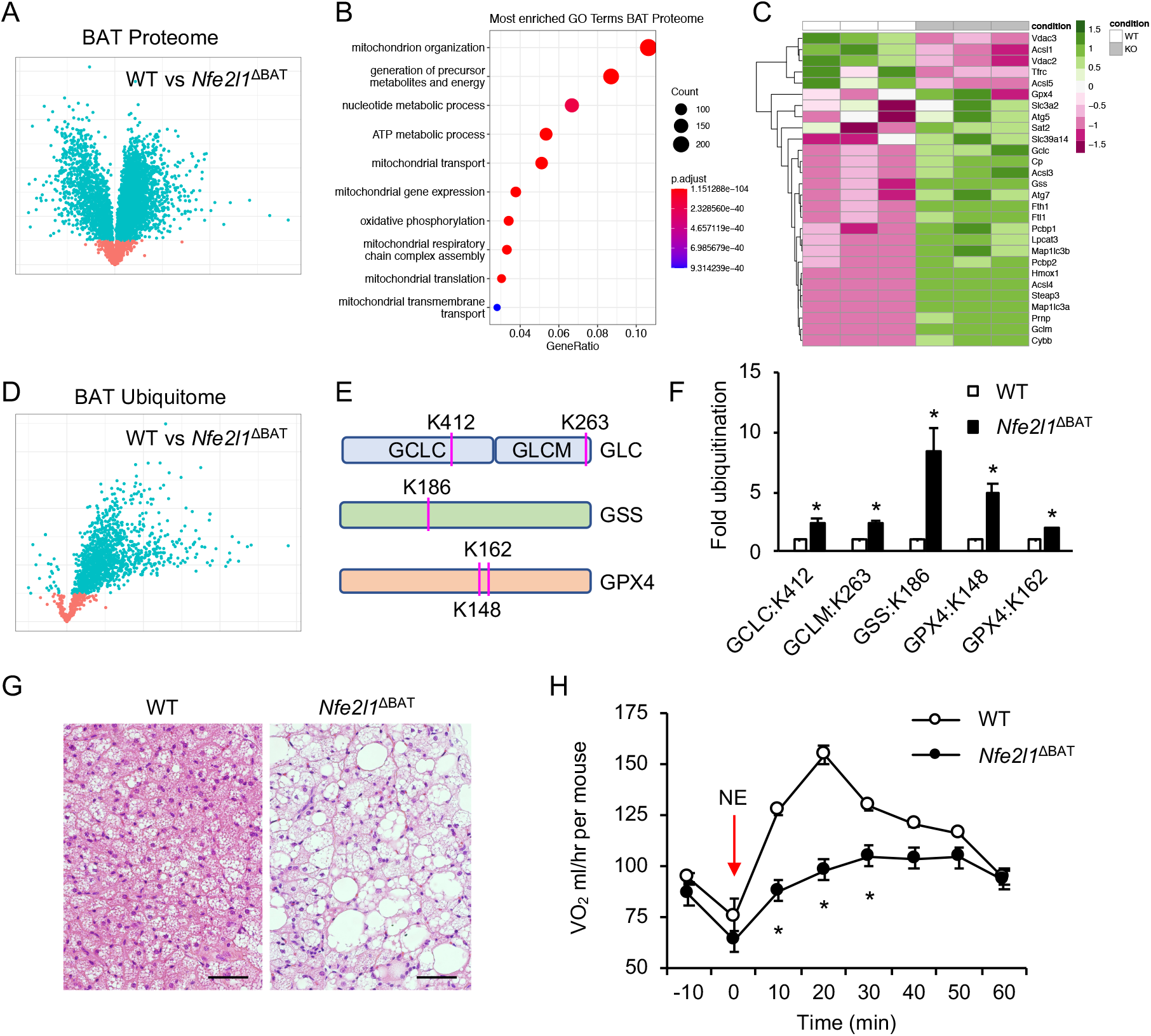
Hallmarks of ferroptosis and hyperubiquitination in Nfe2l1deficiency in vivo. **(A)** Volcano Blot of the BAT proteome from Nfe2l1^ΔBAT^ vs. WT. *P*_adj_ < 0.1 indicated by blue color (*n* = 3 biol. replicates) **(B)** Top 10 enriched gene ontology (GO) terms in the proteome of BAT from Nfe2l1^ΔBAT^ vs. WT (*n* = 3 biol. replicates) **(C)** Heatmap representing the significantly altered expressions of proteins of the KEGG pathway Ferroptosis (*n* = 3 biol. replicates) **(D)** Volcano Blot of the BAT ubiquitome from Nfe2l1^ΔBAT^ vs. WT. *P*_adj_ < 0.1 indicated by blue color (*n* = 3 biol. replicates) **(E,F)** (E) Localization and (F) fold change of ubiquitination of GSH regulating enzymes Gclc, Gclm and Gss as well as Gpx4 in BAT ubiquitome from Nfe2l1^ΔBAT^ vs. WT (*n* = 3 biol. replicates). **(G)** Histology of brown adipose tissue (BAT) from brown adipocyte-specific Nfe2l1-deficient mice (*Nfe2l1*^ΔBAT^) and wild-type (WT) controls. Scale bar: 50 μM **(H)** Oxygen consumption after norepinephrine (NE) injection in *Nfe2l1*^ΔBAT^ and WT controls (*n* = 4 biol. replicates). Statistical significance: \**P*_adj_ < 0.05, by student’s T-test **(F, H)**

## 4. Discussion

Promising discoveries have raised hope for clinical applications of ferroptosis targeting agents, but compared to other cell death modalities, ferroptosis still remains mechanistically insufficiently understood. The combination of specific iron-dependent, lipid-mediated oxidative stress in the context of ferroptosis is potentially linked to proteotoxicity, both through oxidative protein damage by peroxides as well as by maladaptive GPX4 activity. While the relevance of the UPS system in maintaining proteostasis is well known, its role in ferroptosis is unclear. Here we find that proteasomal function and its adaptive regulation by NFE2L1 is a novel mechanism that specifically protects cells from ferroptosis.

Our data show that both pharmacological and genetic induction of ferroptosis is associated with a dynamic depletion and partial restoration of proteasome abundance and activity. By itself this is a remarkable und unprecedented finding, as usually proteasome activity is only modulated moderately and examples of highly dynamic regulation of proteasome function are rare [18]. One could argue that the reduced proteasomal activity is only secondary to the cell death, but proteasomal inhibition already occurs already in the early phase of ferroptosis while cell viability is still unaltered. On the contrary, cells that survive ferroptosis induction show increased proteasomal activity, most likely as a compensatory mechanism due to restoring proteostasis after the proteasomal inhibition in the early phase. Of note, the timing and magnitude of this regulation depends on the nature ferroptosis inducer.

Our data show that this adaptive restoration of proteasome function is mediated by NFE2L1. NFE2L1 is a key transcription factor for increasing proteasome function whenever UPS is challenged [16, 26]. We show that NFE2L1 is activated by both pharmacological and genetic induction of ferroptosis, and this effect was largely independent of increased transcription. Based on the current literature [16, 26], it is likely that the activation of NFE2L1 is directly related to the ferroptosis-induced reduction in proteasomal activity, which causes accumulation of the activated form of NFE2L1, similar to what is seen with proteasome inhibitors. This adaptive activation of NFE2L1 is protective against ferroptosis, as depletion of NFE2L1 reduces proteasomal gene expression and sensitizes cells to ferroptosis.

It is important to point out that cell death induced by proteasome dysfunction [27] is not per se related to ferroptosis. However, treating with proteasome inhibitors sensitizes cells to ferroptosis, which is further enhanced by silencing of Nfe2l1. These results clearly implicate UPS as a key cellular system in the suppression of ferroptosis. Future studies will be required to delineate which aspect of UPS dysfunction are related to ferroptosis. A hallmark of disturbed UPS is ER stress, which has a strong impact on lipid metabolism [28]. Determining the organelle-specific lipidome and oxylipidome will be important to identify lipid species contributing to ferroptosis sensitization. Also, mitochondrial function is tied to UPS [18], representing another potential link of iron homeostasis, oxidative stress, ferroptosis. Finally, a major role of ubiquitin-mediated proteasomal protein degradation is recycling of unwanted, obsolete of damaged proteins.

In that vein, we found that Gpx4 as well as all enzymes required for the synthesis of GSH, the substrate of Gpx4, are hyper-ubiquitinated in Nfe2l1 deficiency in mice. This is in line with previous findings that increased levels of GPX4 ubiquitination sensitize to ferroptosis in vitro [12], but if this relates to changes in the enzymatic activity of GPX4 remains to be investigated. GPX4 has also been shown to not only use GSH as a substrate, but also other protein thiols, especially when GSH is limiting [10]. As shown here, ferroptosis disturbs proteasome function, any putative thiol donors might not be replenished fast enough to contribute to the pool of reducing agents. Likewise, during the initiation of ferroptosis, proteasomal inhibition might reduce substrate availability for GSH synthesis, as proteasomal inhibition causes amino acid shortage [27]. Here, recycling of substrate amino acids for the GPX4 pathway might be a relevant mechanism that requires further investigation. As we found multiple other proteins implicated in ferroptosis, for instance ACSL4 [25] to be impacted by NFE2L1-mediated proteasome function and ubiquitination [18], more research is needed to characterize the biochemical role of these lysine residues for protein degradation and function, respectively.

Somewhat surprisingly, erastin, which inhibits SLC7A11 and therefore limits cellular cystine import and cysteine levels, decreased both NFE2L1 protein levels and activity despite mRNA levels being robustly elevated. This is in stark contrast to the effects of RSL3, FIN56, and genetic induction of ferroptosis. Whereas RSL3 and FIN56 are rather specific, erastin shows broader reactivity, as regulation of SLC7A11, VDAC2/3 and p53 by erastin have been reported [29, 30]. It is therefore possible that the induction of ferroptosis by erastin has other, secondary effects, that mask or counteract the impact on NFE2L1. This finding might further our mechanistic understanding of erastin and NFE2L1 biology.

In search for the potential biological significance of our observation, we turned to brown adipocytes as a model system. These cells display highly adaptive NFE2L1-mediated proteasome function in response to physiological cold stimulation [18]. In the absence of NFE2L1, BAT undergoes a progressive loss of thermogenic capacity associated with aberrant mitochondrial function and altered lipid metabolism [18]. Our data here show that part of this tissue degeneration is likely related to ferroptosis. Further genetic evidence is required to verify the contribution of ferroptosis to thermogenic decline, a scenario that is observed with ageing in humans [31]. Interestingly, BAT is characterized by a unique combination of high iron and lipid content. Indeed, several oxidized lipids have thermogenic effects in BAT [32, 33]. Therefore, it seems reasonable that BAT protects itself from ferroptosis with specialized mechanisms, one being high activity of NFE2L1. Shielding brown adipocytes from ferroptosis might be a potential strategy to prevent ageing-associated decline in BAT function.

On the other hand, proteasomal inhibitors such as bortezomib are already in clinical use in the therapy against multiple myeloma. Clinical trials with the erastin analogue PLRX 9393 (ClinicalTrials.gov Identifier: NCT01695590) were carried out for the same disease but so far remained inconclusive [34]. Our data support the notion that the route of ferroptosis has differential effects on NFE2L1 and UPS, which highlight the need for more studies in this context. Furthermore, we show that bortezomib by itself does not cause ferroptotic cell death, but careful titration of proteasome inhibition accelerates ferroptosis in human patient-derived GPX4-mutated cells. This finding may open possibilities for combined therapy with inducers of ferroptosis [34] and proteasomal inhibition, as they do not only have additional toxicity but rather synergistic potential.

Finally, our study enhances the understanding of the pathogenesis underlying SSMD, as we identify NFE2L1-mediated proteasomal function as protective mechanism against ferroptosis in patient fibroblasts. Obviously, translating this basic finding to better treatment options is challenging, yet NFE2L1 might be a feasible therapeutic target worthy of further investigations, either by gene therapy or pharmacological activation. Dietary substitution to replenish the amino acid pool used for GSH synthesis could be a simple alternative approach to combat the effects of proteasomal downregulation in the patients’ cells.

## 5. Conclusions

Our study highlights the protective nature of the NFE2L1-proteasome pathway against ferroptosis. Manipulation of NFE2L1 activity might enhance ferroptosis-targeting cancer therapies as well as protect from aberrant ferroptosis in neurodegeneration, metabolism, and beyond.

## Acknowledgement

We thank Julia Schluckebier, Silvia Weidner, Thomas Pitsch, Jonas Wanninger, and Maude Giroud for excellent technical assistance. We thank all lab members for discussions and the enjoyable atmosphere. We thank Scott Dixon and Giovanni Forcina for discussions and scholarship. We thank CureGPX4.org, a non-profit organization to create treatments of patients affected by diseases caused by mutations in GPX4 gene, for sharing the patient-derived and control fibroblasts. We thank Brice Emanuelli for providing us with WT1 cells and Siegfried Ussar for the imBAT cells. We thank Silke Meiners and colleagues for help with the native PAGE. S.K. was supported by the LMU Medical Faculty program FöFoLe for MD students. Work in the Conrad lab is supported by the Deutsche Forschungsgemeinschaft (DFG) CO 291/7-1, the DFG Priority Program 2306 (CO 297/9-1; CO 297/10-1), the Ministry of Science and Higher Education of the Russian Federation (075-15-2019-1933) and the Else Kröner-Fresenius-Stiftung (2020_EKTP19). M.C. is further funded by the European Research Council (ERC) under the European Union’s Horizon 2020 research and innovation programme (grant agreement No. GA 884754). A.B. was supported by the DFG Priority Program 2306 (BA 4925/2-1), the Deutsches Zentrum für Herz-Kreislauf-Forschung DZHK Junior Research Group Program (JRG), and the ERC Starting Grant PROTEOFIT. The graphical abstract was created using BioRender.com. We apologize to colleagues whose work we could not cite due to space limitations.

## Competing interest

The authors declare no competing financial interests related to this work.

## Author contributions

S.K. designed and performed experiments, analyzed data, and wrote the manuscript. A.J., N.W., A.O., and M.C. performed experiments and analyzed data. A.B. designed and performed experiments, analyzed data, and wrote the manuscript. All authors read and commented on the manuscript.

## Abbreviations

ACSL4: Acyl-CoA Synthetase Long Chain Family Member 4
BAT: Brown adipose tissue
ER: endoplasmic reticulum
GCL: Glutamate-cysteine ligase
GPX4: Glutathione peroxidase-4
GSH: glutathione
GSS: Glutathione synthetase
NFE2L1: Nuclear factor erythroid-2, like-1
SSMD: Sedaghatian-type Spondylometaphyseal Dysplasia
UPS: ubiquitin-proteasome system

---

**Supplementary Table 1:**
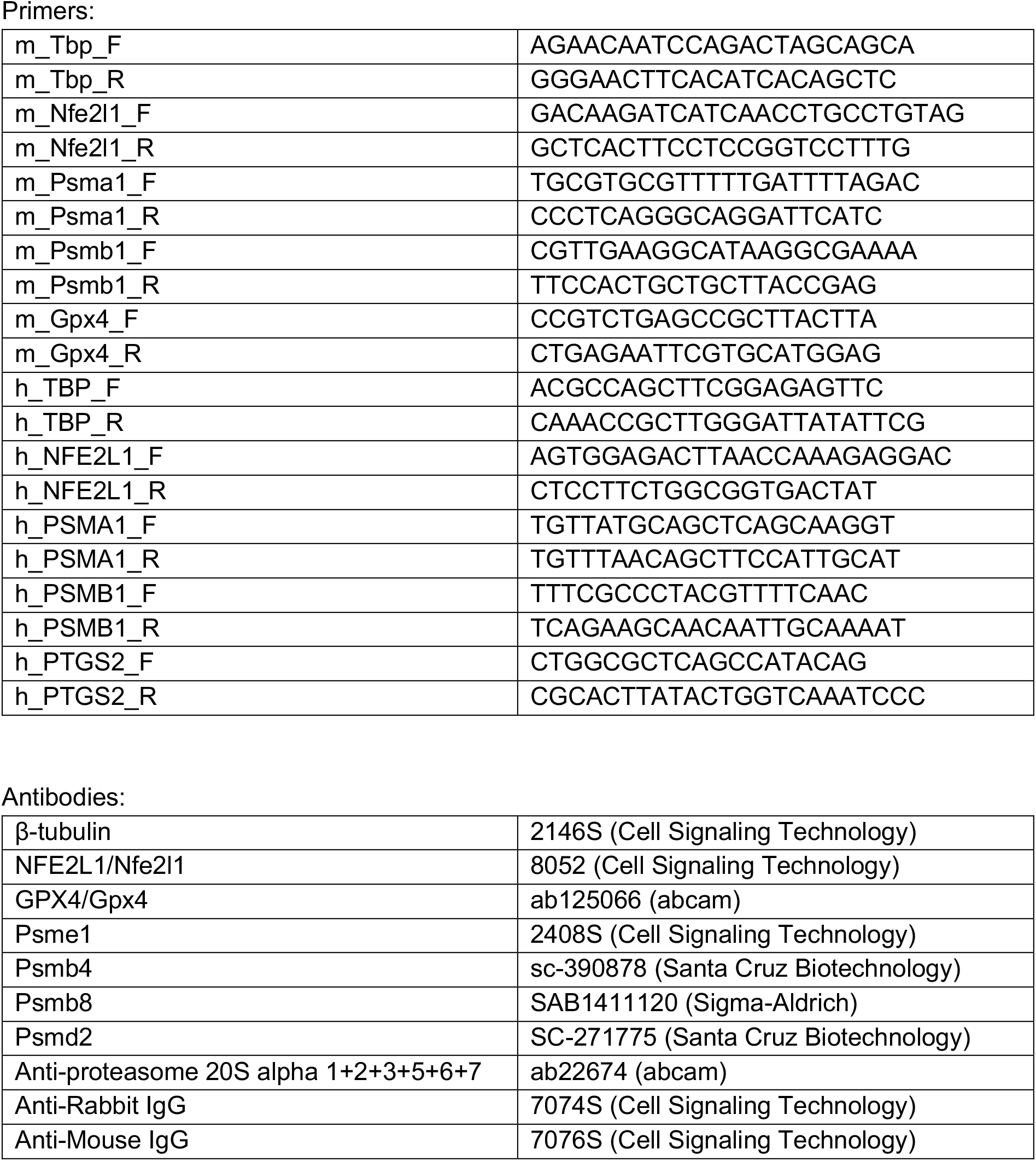

## Notes

### Competing Interest Statement

The authors have declared no competing interest.

